# From static to temporal network theory – applications to functional brain connectivity

**DOI:** 10.1101/096461

**Authors:** William Hedley Thompson, Per Brantefors, Peter Fransson

**Affiliations:** Department of Clinical Neuroscience, Karolinska Institutet, Stockholm, Sweden

**Keywords:** resting-state, temporal network theory, temporal networks, functional connectome, dynamic functional connectivity

## Abstract

Network neuroscience has become an established paradigm to tackle questions related to the functional and structural connectome of the brain. Recently, there has been a growing interest to examine the temporal dynamics of the brain's network activity. While different approaches to capture fluctuations in brain connectivity have been proposed, there have been few attempts to quantify these fluctuations using temporal network theory. Temporal network theory is an extension of network theory that has been successfully applied to the modeling of dynamic processes in economics, social sciences and engineering. The objective of this paper is twofold: (i) to present a detailed description of the central tenets and outline measures from temporal network theory; (ii) apply these measures to a resting-state fMRI dataset to illustrate their utility. Further, we discuss the interpretation of temporal network theory in the context of the dynamic functional brain connectome. All the temporal network measures and plotting functions described in this paper are freely available as a python package *Teneto*.

## Introduction

It is well-known that the brain’s large-scale activity is organized into networks. The underlying organization of brain’s infrastructure into networks, at different spatial levels, has been dubbed the brain’s functional and structural connectome (1, 2). Functional connectivity, derived by correlating the brain’s activity over a period of time, has been successfully applied in both fMRI (3, 4, 5, 6) and MEG (7, 8, 9), yielding knowledge about functional network properties (10, 11, 12, 13) which has been applied to clinical populations (14, 15).

In parallel to research on the brain’s connectome, there has been a focus on studying the dynamics of brain activity. When the brain is modeled as a dynamic system, a diverse range of properties can be explored, prominent examples of this are metastability (16, 17, 18, 19, 20) and oscillations (21, 22, 23). Brain oscillations, inherently dynamic, have become a vital ingredient in proposed mechanisms ranging from psychological processes such as memory (24, 25, 26), attention (27, 28), to basic neural communication of a top-down and bottom-up type of information transfer (29, 30, 31, 32, 33, 34, 35, 36).

Recently, approaches to study brain connectomics and the dynamics of neuronal communication have started to merge. A significant amount of work has recently been carried out that aims to quantify dynamic fluctuations of network activity in the brain using fMRI (37, 38, 39, 40, 41, 42) as well as MEG (7, 43, 9, 44, 35). This research area to unify brain connectommics with the dynamic properties of neuronal communication has been called the “dynome” (45) and the “chronnectome” (46). As the brain can quickly fluctuate between different tasks, the overarching aim of this area of research is to understand the dynamic interplay of the brain’s networks. The intent of this research is that it will yield insight about the complex and dynamic cognitive human abilities.

Although temporal network theory has been successfully applied in others fields, (e.g. social sciences), its implementation in network neuroscience is limited. In this paper, we first provide an introduction to temporal network theory by extending the definitions and logic of static network theory. Thereafter, we define temporal network measures. Finally, we apply these measures to a resting-state fMRI dataset acquired during eyes open and eyes closed conditions, revealing differences in dynamic brain connectivity between conditions.

### From static networks to temporal networks

We begin the introduction to temporal network theory by expanding upon the definitions of network theory. In network theory, a graph (*G*) is defined as a set of nodes and edges:

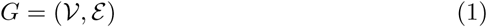

𝓥 is a set containing *N* number of nodes. *ɛ* is a set of tuples that represents edges or connections between each pair of nodes (*i,j*): *i,j* ∈ 𝓥. The graph may have binary edges (i.e. an edge is either present or absent) or it may be weighted, often normalized between 0 and 1, to represent the magnitude of connectivity. When each edge has a weight, the definition of *ɛ* is extended to a 3-tuple (*i,j, w*) where *w* denotes the weight of the edge between *i* and *j*.

*ɛ* is often represented as a matrix of the tuples, which is called a connectivity matrix, *A* (sometimes the term “adjacency matrix” is used). An element of the connectivity matrix *A_i,j_* represents the degree of connectivity for a given edge. When *G* is binary, *A_i,j_* = {0,1} and in the weighted version *A_i,j_* = *w*. In the case of *A_i,j_* = *A_j,i_*,for all *i* and *j*, the matrix is considered undirected and, when this is not the case, directed.

One appeal of network theory is the range of topics it can encompass. A set of nodes can be a group of people, a collection of cities, or an ensemble of brain regions. Each element in the nodal set can represent vastly different things in the world (e.g. a person: Ashely; a city: Gothenburg; or a brain region: the left thalamus). Likewise, edges can represent a range of different types of connections between their respective nodes (e.g. friendship, transportation or neural communication). Regardless of what the network is modeling, many different properties regarding the patterns of connectivity between nodes can be quantified, examples being centrality measures, hub detection, small world properties, clustering and efficiency (see 47, 2, 48 for detailed discussions).

It is important to keep in mind that a graph is only a representation of some state of the world being modeled. The correspondence between the graph model and the state of the world may decrease due to aggregations, simplifications and generalizations. Adding more information to the way nodes are connected can entail that *G* provides a better representation, thus increasing the likelihood that subsequently derived properties of the graph correspond with the state of the world being modeled. One simplification made in eq. 1 is that two nodes can be connected by one edge only. Such a simplification may not be appropriate for all questions. For example, if we wanted to model multiple transportation links between several cities (e.g. rail, road, and air), we will need multiple types of edges if these different transportation links are of importance. In some circumstances, the type of link is irrelevant. For example, when studying the spread of a disease, the only relevant information needed might be that people can move between cities. However, if we are managing shipping routes where different transportation routes correspond to different prices or shipping time, then this might be important information. Similarly, networks (e.g. social networks) develop and change over time. In social networks, friendships are started and can end.

To capture such additional information in the graph, edges need to be expressed along additional non-nodal dimensions. We modify eq. 1 to:

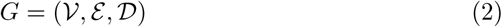

where 𝓓 is a set of the additional non-nodal dimension. In the case of multiple additional dimensions, 𝓓 is a class of disjoint sets where each dimension is a set. Eq. 2 is sometimes referred to as a *multigraph.* For example, 𝓓 could be a set containing three transportation types {“rail”, “road”, “air”} or a temporal index {“2014”,“2015”,“2016”}.

In a static graph, the edges *ɛ* are elements that contains pairs (2-tuples) and thus represent connections between two nodes. In a multigraph, *ɛ* consists of (|𝓓| + 2)-tuples for binary graphs, where |𝓓| expresses the number of sets in 𝓓. Here, the addition of two is because of the original two node indexes, which defines the edge in eq. 1. If time is the only dimension in 𝓓, then an element in *ɛ* is a triplet: (*i,j,t*): *i,j* ∈ 𝒱,*t* ∈ *D*. When *G* is weighted, *ɛ* contains (2 × |𝓓|) + 2)-tuples as *w* becomes the size of 𝓓, representing one weight per edge.

While some of the measures presented here are valid for all types of multigraphs, time is a special case since it comprises of an ordered set. There is no loss of information if {“rail”, “road”, “air”} becomes {“road”,“air”,“rail”}. However, when 𝓓 contains temporal information, the order is crucial. Thus, a temporal network is when a multigraph has an ordered set in 𝓓 that represents time.

From the description above, it is evident that multiple sets can be included in a multigraph. For example, consider a set of three cities. Let 𝓓 be ({“rail”,“road”,“air”},{“2014”,“2015”,“2016”}). Here, each edge is expressed in a multigraph by a 4-tuple which indexes the two cities, transportation type, and year. Analogously, it is conceivable that a detailed network description of the human connectome may include information regarding an edge’s presence in time, frequency and task context. Thus, a multigraph representation of the connectome may, for example, include information on edges that are conditional to (i) a delay of 100ms after stimulus onset (time), (ii) presence of gamma oscillations (frequency), and (iii) the presence of an n-back exercise (task context). In such complex multigraphs, the temporal network measures presented in this paper can be used to examine relationships across time, but it will require fixing or aggregating over the other dimensions. However, more complex measures that consider all non-nodal dimensions have been proposed elsewhere (e.g. 49).

For the remaining parts of this paper we will only consider the case when 𝓓 contains an ordered set of temporal indexes and when *G* is binary and undirected. In this case, each edge is indexed by *i,j* and *t*. To facilitate readability, connectivity matrices are written as 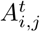 i.e. with the temporal index of 𝓓 in the superscript.

We can illustrate the concept of temporal networks by a simple example that models the evolution of friendship between three individuals (Ashley, Blake and Casey). A friendship between two of them is represented by an edge (Figure 1A). We then add an additional dimension, time, to the graph which includes the following temporal indexes: 𝓓 ={2014,2015,2016}. Let us now presume that in 2014, only Ashley and Blake knew each other. In 2015, Ashley became friends with Casey. In 2016, Blake and Casey also became friends. The temporal evolution of friendships among a small social network can be projected onto a slice graph representation (Figure 1B). Here, it becomes apparent that Casey is the person in this small social network that shows the largest change in friendship over time, a property which cannot be depicted in a static graph.

**Figure 1:**
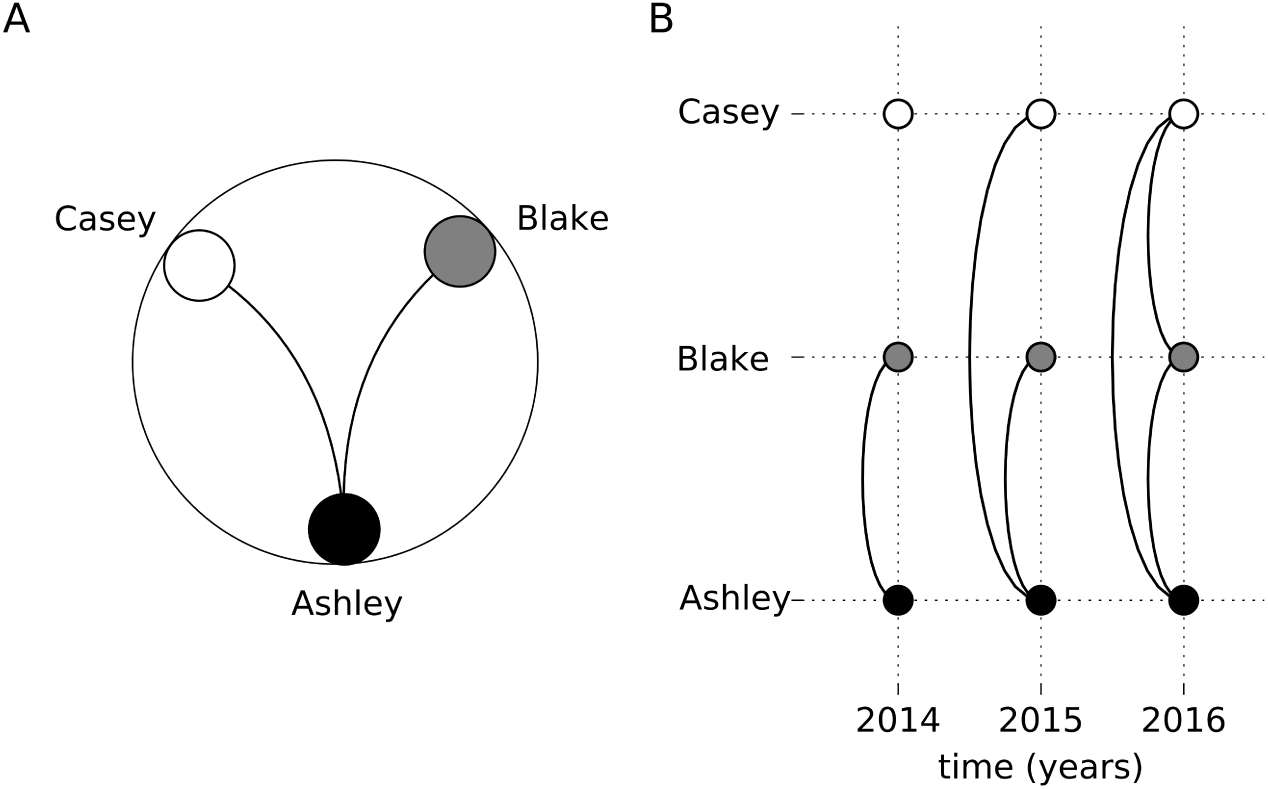
A slice graph example of friendship in a small social network evolving over three years. Each person (node) is a row with a circle at each year (the non-nodal dimension). Lines display represent friendship (edges) between the nodes.

The small social network in Figure 1B shows snapshots of connectivity through time. At each time point, there is a connectivity matrix which is sometimes called a *graphlet* (50, 40, 41). A graphlet is a complete two dimensional connectivity matrix, akin to static networks, but each graphlet is only a part of the entire temporal network. It useful to describe what type of graphlet that is being used. For example we call a graphlet expressing time for a time-graphlet or t-graphlet (41) . Likewise, a graphlet expressing frequency information has been called a frequency graphlet or f-graphlet (40).

Instead of representing the data with multiple graphlets, **ɛ** can be used to derive the *contact sequence* containing the nodes and temporal index (51). Unlike the graphlet representations, which must be discrete, contact sequences can also be used on continuous data and, when connections are sparse, a more memory efficient way to store the data.

## Measures for temporal networks

Once the t-graphlets have been derived, various measures can be implemented in order to quantify the degree and characteristics of the temporal flow of information through the network. We begin by introducing two concepts which are used in several of the temporal network measures that will be defined later. The focus is on measures that derive temporal properties at a local level (i.e. per node or edge) or a global level (see Discussion for other approaches). We have limited our scope to describe only the case of binary, undirected and discrete t-graphlets, although many measures can be extended to continuous time, directed edges, and non-binary data.

### Concept: Shortest temporal path

In static networks, the shortest path is the minimum number of edges (or sum of edge weights) that it takes for a node to reach another node. In temporal networks, a similar measure can be derived. Within temporal networks, we can quantify the time taken for one node to reach another node. This is sometimes called the “shortest temporal distance” or “waiting time”.

Consider the temporal network shown in Figure 2A. Starting at time point 1, the shortest temporal path for node 1 to reach node 5 is 5 time units (Figure 2B, red line). Here, only one edge was traveled per time point. However, deriving the shortest temporal path becomes a more complex task when considering that multiple edges can be traveled per time point. For example, node 5 at time 2 can reach node 3 in one time step, if multiple edges are allowed to be traveled (Figure 2C, red line). If multiple edges cannot be traveled, then the shortest path for node 5 to reach node 2, starting at time point 2, is 5 time units (Figure 2C, blue line).

**Figure 2:**
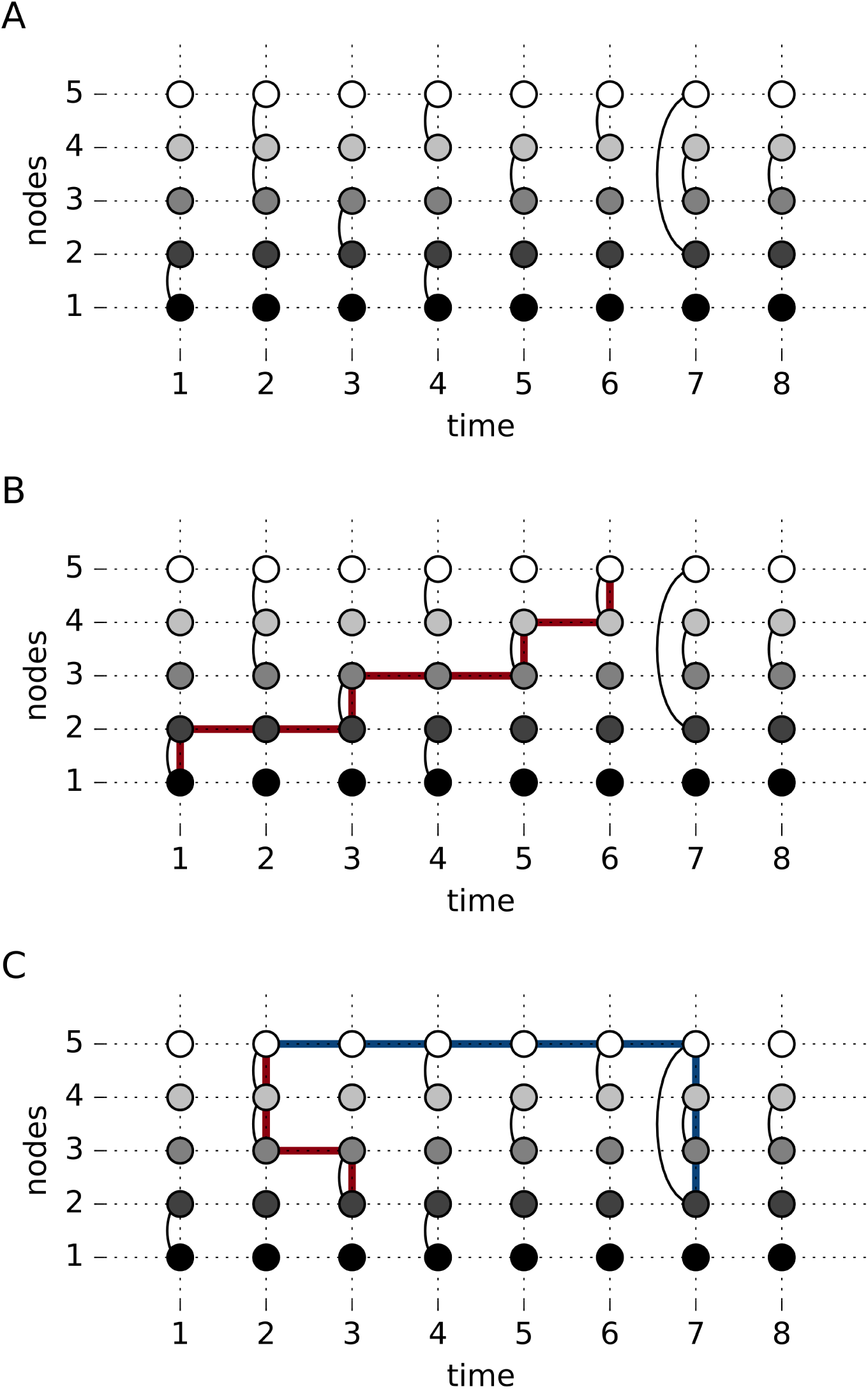
Illustration of the concept of shortest temporal path. (A) The basic layout of a temporal network viewed in a slice graph representation. (B) the red line indicates the shortest temporal path possible for node 1 to reach node 5. (C) shows the difference in shortest paths that arise when allowing for a single (blue line) or multiple (red line) edges to be traveled at a single time point.

When calculating the shortest temporal path, a parameter must be set that restrains how many edges per time point that can be traveled. This parameter will depend on the temporal resolution of the data and is best chosen given previously established knowledge of the dynamics of the data. For fMRI, where the temporal resolution is in seconds, it makes sense to assume that several edges can be traveled per unit of time. Contrarily, in MEG where the resolution are in the range of milliseconds, it is more reasonable to place a limit on the number of edges that can be traveled per time unit.

Regarding the shortest temporal path, it is useful to keep in mind that the path is rarely symmetric, not even so when the t-graphlets themselves are symmetric. This is illustrated by considering the network shown in Figure 2A for which it takes 5 units of time for node 1 to reach node 5 when starting at *t =* 1. However, for the reversed path, it only takes 3 units of time for node 5 to reach node 1 (allowing for multiple edges to be traveled per time point).

### Concept: Inter-contact time

The inter-contact time between two nodes is defined as the temporal difference for two consecutive non-zero edges between those nodes. This definition differs from the shortest temporal paths in so far as it only considers direct connections between two nodes. Considering Figure 2A, the inter-contact times between nodes 4 and 5 become a list [2,2] as there are edges present at time points 2, 4 and 6. Each edge will have a list of inter-contact times. The number of inter-contact times in each list will be one minus the amount non-zero edges between the nodes. Unlike the shortest temporal paths, graphs that contain inter-contact times will always be symmetric.

### Nodal measure: Temporal centrality

Akin to degree centrality in the static case, where the sum of edges for a node is calculated, a node’s influence in a temporal network can be calculated in a similar way. The difference from its static counterpart is that we additionally sum the number of edges across time. Formally, temporal degree centrality, *D^T^*, for a node *i* is computed as

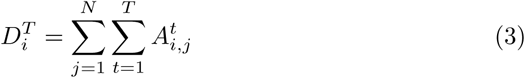

where *T* is the number of time points, *N* is the number of nodes and 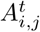 is a graphlet.

While providing an estimate of how active or central a node is in a temporal network, temporal degree centrality does not quantify the temporal order of the edges. This is illustrated in Figure 3, where node 3 and node 2 have identical temporal degree centrality despite having very different temporal ordering of their edges.

**Figure 3:**
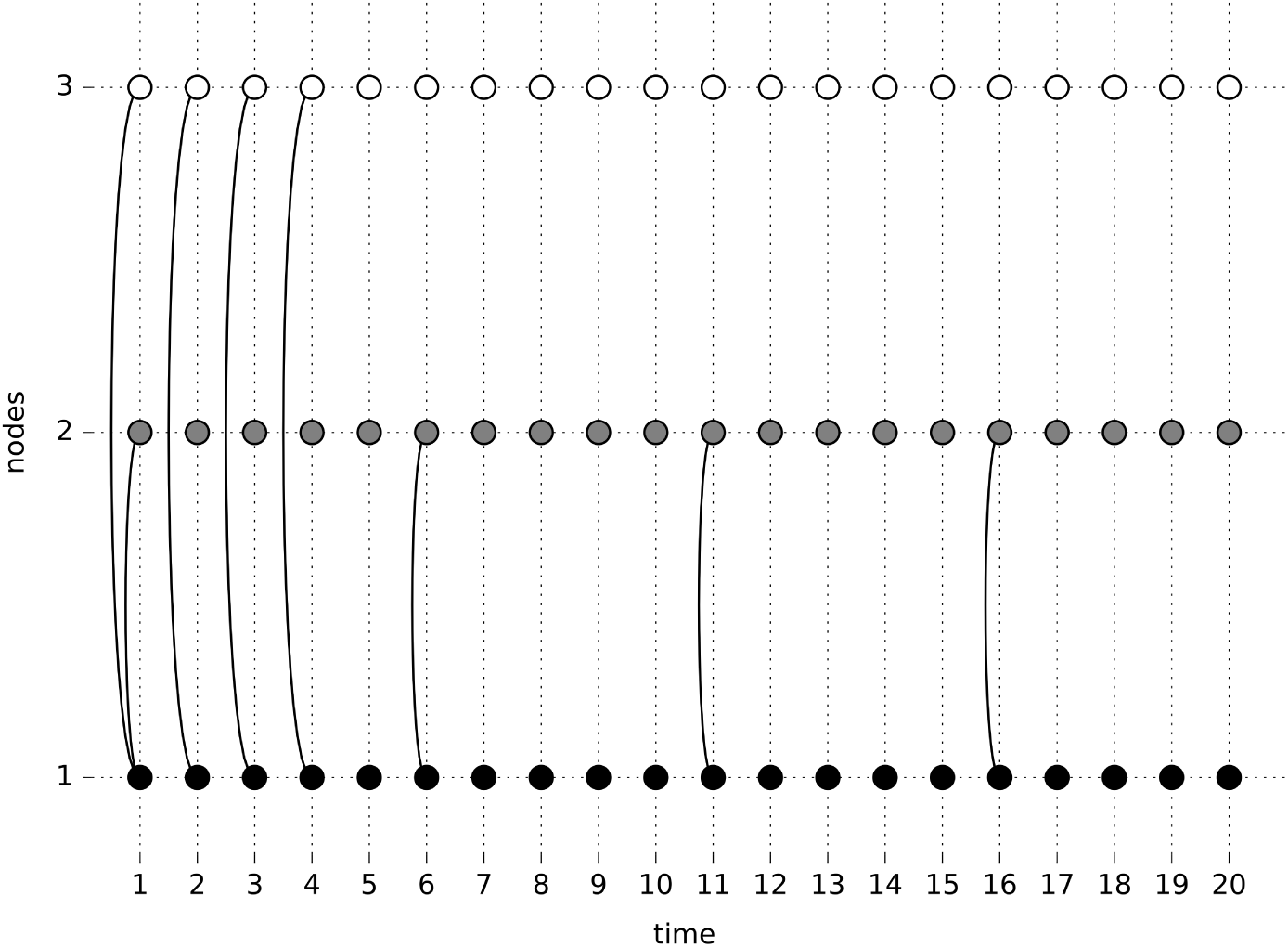
A slice graph representation of a simple example of a temporal network that illustrates the conceptual difference between temporal degree centrality and temporal closeness centrality.

### Nodal measure: temporal closeness centrality

A centrality measure that considers the temporal order is temporal closeness centrality (52). This is an extension of closeness centrality which is the inverse sum of the shortest temporal. Temporal closeness centrality is calculated by the average of the shortest temporal paths between two nodes *d^τ^* as

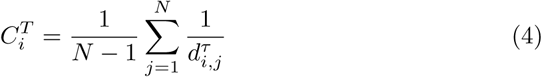

where 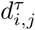 is the average shortest path between nodes *i* and *j*. Like its static counterpart, if a node has shorter temporal paths compared to other nodes, it will have a larger temporal closeness centrality.

Consider the example given in Figure 3 that shows a temporal sequence of connectivity among three nodes over 20 time points. Note that the temporal degree centrality is identical for both node 2 and node 3, while degree centrality for node 1 is twice as large. Node 2 has the largest temporal closeness centrality since the time between edges for node 2 are longer than for node 3, which has the lowest value of temporal closeness centrality.

### Edge measure: Bursts

Bursts have been identified using temporal network theory as an important property for many processes in nature (53, 54, 55, 56, 57). A hallmark of a bursty edge is the presence of multiple edges with short inter-contact times, followed by longer and varying inter-contact times. In statistical terms such a process is characterized with a heavy tailed distribution of inter-contact time probabilities. Numerous patterns of social communication and behaviour have been successfully modeled as bursty in temporal network theory, including email communication (58, 53), mobile phone communication (59), spreading of sexually transmitted diseases (60), soliciting online prostitution (61), and epidemics (62). With regard to network neuroscience, we have recently shown that bursts of brain connectivity can be detected in resting-state fMRI data (41). Furthermore, bursty temporal patterns have also been identified for the amplitude of the EEG alpha signal (63, 64, 65).

There are several strategies to quantify bursts. A first indication of whether a time series of brain connectivity between two nodes is bursty or not is simply to plot the distribution of inter-contact times. Thus, the complete distribution of *τ* for a given edge contains information of the temporal evolution of brain connectivity. However, there are other methods available to quantify bursts. One example is the burstiness coefficient (*B*) first presented in ref. (66) and formulated for discrete graphs in ref. (51):

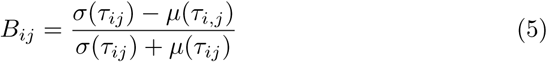

where *τ_ij_* is a vector of inter-contact times between nodes *i* and *j* through time. When *B* > 0, it is an indication that the temporal connectivity is bursty. This occurs when the standard deviation *σ*(*τ*) is greater than the mean *μ*(*τ*). In eq. 5, bursts are calculated per edge which can be problematic when having limited data. Functional imaging sessions must be long enough in order to be able to accurately establish whether a given temporal distribution is bursty or not (too few inter-contact times will entail a too poor estimation of *σ* to accurately estimate *B*). Typically, for resting state fMRI datasets acquired during rather short time spans (5-6 minutes) with low temporal resolution (typically 2-3 seconds), it might be difficult to quantify *B* in a single subject. A potential remedy in some situations is to compute *B* after concatenating inter-contact times across subjects.

Eq. 5 calculates the number of bursts per edge. This can easily be extended to a nodal measure by summing over the bursty coefficients across all edges for a given node. Alternatively, a nodal form of *B* can be calculated by using all the inter-contact times for all j, instead of averaging over *j* in *B_ij_*. Finally, if a process is known to be bursty, instead of quantifying *B*, it is possible to count the number of bursts present in a time series.

### Global measure: Fluctuability

While centrality measures provide information about the degree of temporal connectivity and bursts describe the distribution of the temporal patterns of connectivity at a nodal level, one might additionally want to retrieve information about the global state of a temporal network. To this end, fluctuability aims to quantify the temporal variability of connectivity. We define fluctuability *F* as the ratio of the number of present edges in *A* over the grand sum of *A_t_*

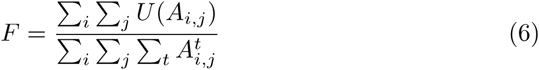

where *U* is a function that delivers a binary output as follows: *U(A_i,j_)* is set to 1 if at least one of an edge occurs between nodes *i* and *j* across time *t* = 1, 2,…, *T*. If not, *U*(*A_i,j_)* is set to zero. This can be expressed mathematically as:

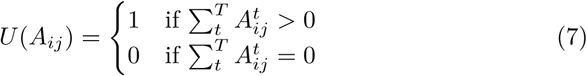

where *T* is the total number of time points and *A* has at least one non-zero edge. From the definition given in eq. 6 it follows that the maximum value of *F* is 1 and this value only occurs when every edge is unique and occurs only once in time.

While the above definition of fluctuability may seem counter-intuitive, it is an adequate measure to quantify the temporal diversity of edges. *F* reveals how connectivity patterns within the network fluctuates across time. To see this, consider the networks shown in Figures 4A and 4B, for which there are two edges present at each time point. There are only three different unique edges in Figure 4A, meaning that the sum of *U* is 3 for the network shown in Figure 4A. However, there is a greater fluctuation in edge configuration for the network shown in Figure 4B and all six possible edges are present (entailing that the sum of *U* is equal 6). Since both networks have in total 24 connections over time, it becomes easy to see that the network shown in Figure 4B has a twice as large value of *F* compared to the network shown in Figure 4A.

**Figure 4:**
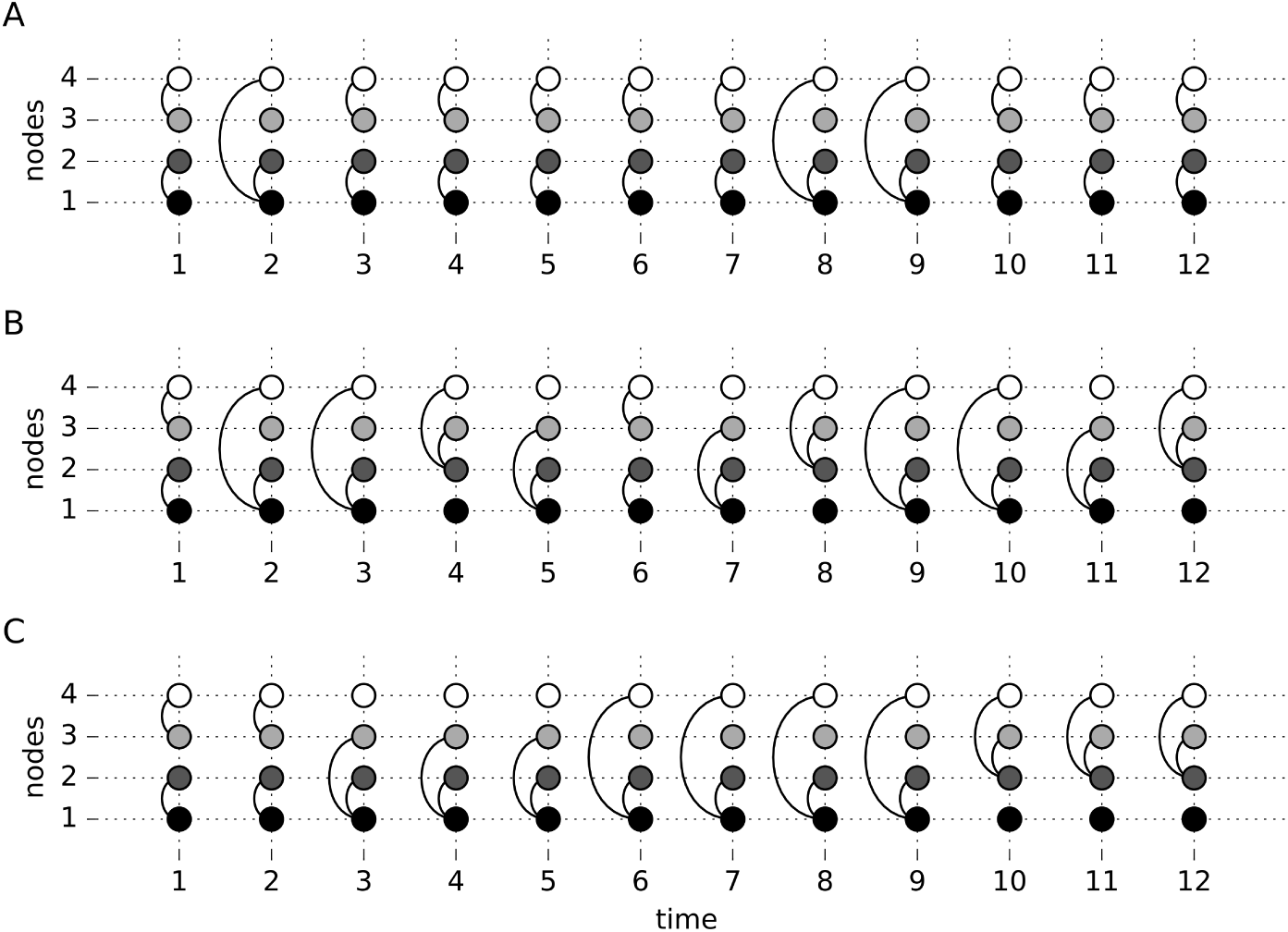
An illustration of the fluctuability and volatility measures. The temporal networks shown in panels A, B and C all have the same number of nodes and edges but they differ in fluctuability and volatility. (A) has low fluctuability (*F* = 0.125) and volatility (*V* = 0.73). (B) The network with the highest volatility of all three (*V*=2.55) networks and equal fluctuability (*F*=0.25) compared to the network in panel C. (C) A network with lower volatility than B (*V* = 1.27) but equal fluctuability (*F*=0.25).

Notably, fluctuability is insensitive to the temporal order of connectivity. For example, the networks depicted in Figures 4B and 4C have the same fluctuability, despite having a considerably different temporal orders of edge connectivity. Thus, fluctuability can be used as an indicator of the overall degree of spatial diversity of connectivity over time.

The definition of fluctuability can be changed to work on a nodal level. To achieve this, the summation in eq. 6 is applied over only one of the nodal dimensions. Note that for nodes with no connections at all, the denominator will be 0 and, to circumvent this hindrance, nodal fluctuability 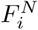 is defined as:

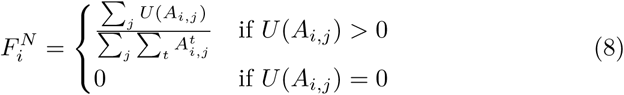

### Global measure: Volatility

One possible global measure considering the temporal order is to quantify how much, on average, connectivity between consecutive t-graphlets changes. This indicates how volatile the temporal network is over time. Thus, volatility (*V*) can be defined as:

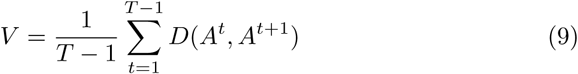

where *D* is a distance function and *T* is the total number of time points. The distance function quantifies the difference between a graphlet at *t* with the graphlet at *t*+1. In all the following examples in this paper for volatility use Hamming distance as it is appropriate for binary data.

Whereas there was no difference in fluctuability between the networks shown in Figures 4B and 4C, there was a difference in volatility, as the network in Figure 4B has more abrupt changes in connectivity compared to the network shown in Figure 4C.

Extensions of the volatility measure are possible. Similar to fluctuability, volatility can be defined at a local level. A per edge version of volatility can be formulated as:

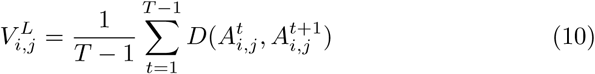

Additionally, taking the mean over *j* in 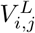 would give an estimate of volatility centrality.

### Global measure: Reachability Latency

Measures of reachability focus on estimating the time taken to “reach” nodes in a temporal network. In ref. 67, both *reachability ratio* and *reachability time* are used. The reachability ratio is the percentage of edges that have a temporal path connecting them. The reachability time is the average length of all temporal paths. However, when applying reachability to the brain, the two aforementioned measures are not ideal given the non-controversial assumption that any region in the brain, given sufficient time, can reach all other regions.

With this assumption in mind, we define a measure of reachability, *reachability latency* that quantifies the average time it takes for a temporal network to reach an a priori defined reachability ratio. This is defined as:

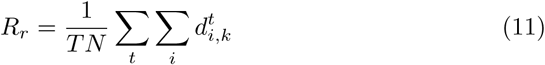

where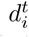 is an ordered vector of length *N* of the shortest temporal paths for node *i* at time point *t*. The value *k* represents the 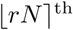 element in the vector, which is the rounded product of the fraction of nodes that can be reached, *r*, with *N* being the total number of nodes in the network.

In the case of *r* =1, (i.e. 100% of nodes are reached), eq. 11 can be rewritten as:

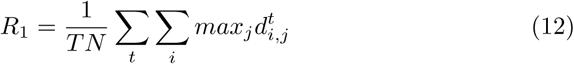

Eq. 12 has been referred to as the *temporal diameter* of the network (68). If eq. 12 is modified and calculated per node instead of averaging over nodes, it would be a temporal extension of node eccentricity.

Unless all nodes are reached at the last time point in the sequence of recorded data, there will be a percentage of time points from which all nodes can not be reached. This effectively reduces their value of *R* as 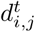 cannot be calculated by *R* is still normalized by *T*. If this penalization is considered too unfair, it is possible to normalize *R* by replacing *T* with *T*\*, which is the number of time points where 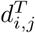 has a real value.

### Global measure: Temporal Efficiency

A similar concept is the idea of temporal efficiency. In the static case, efficiency is computed as the inverse of the average shortest path of all nodes. Temporal efficiency is first calculated at each time point as the inverse average shortest path length for all nodes. Subsequently, the inverse of average shortest path lengths are averaged across time points to obtain an estimate of global temporal efficiency, which is defined as

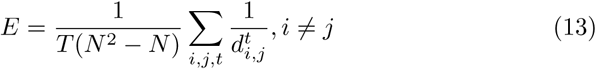

Although reachability and efficiency estimate similar temporal properties, since both are based on the shortest temporal paths, the global temporal efficiency may result in different results than the reachability latency. This is because efficiency is proportional to the average shortest temporal path and reachability to the longest shortest temporal path to reach *r* percent of the network. Similar to the case of static graphs, temporal efficiency can also be calculated on a nodal as well as a global level.

### Summary of temporal network measures

In Table 1 we provide a brief summary of the temporal network measures outlined here, accompanied with a short description. We also signify which measures that are sensitive to temporal order.

### Statistical considerations of temporal network measures

When implementing temporal graph measures, it is important to perform adequate statistical tests in order to infer differences between subject groups, task conditions, or chance. For group comparisons, non-parametric permutation methods are advantageous where the group assignment of the calculated measure can be shuffled between the groups and a null-distribution can be created.

Alternatively, to justify that a measure is significantly present above chance levels, construction of null graphs is needed. There exist multiple ways to create temporal null graphs and they each have their own benefits and drawbacks. One method is to permute the temporal order of entire time series, but this will destroy any auto-correlation present in the data. Another alternative is to permute the phase of the time series prior to thresholding the t-graphlets. A third option would be to permute blocks of time series data. We refer the reader to ref. (51) for a full account of approaches on how to perform statistical tests on measures derived from temporal network theory.

**Table 1:** A summary of the temporal network measures outlined in this article.

| Measure | Description | Order de-<br>pendent | Variable |
| --- | --- | --- | --- |
| Temporal centrality | Number of overall overall connections in time | N | $D^T$ |
| Closeness centrality | The time between connections | Y | $C^T$ |
| Burstiness | the distribution of subsequent connections | Y | $B_{i,j}$ |
| Fluctuability | Ratio of unique edges vs all edges. | N | $F$ |
| Volatility | The rate of change of the graphlets per time-point. | Y | $V$ |
| Reachability latency | Time taken for all nodes to be able to reach each other | Y | $R_r$ |
| Temporal efficiency | Inverse average shortest temporal path. | Y | $E$ |

## Applying temporal network measures onto fMRI data

In the second part of the paper we now turn our attention to the application and interpretation of temporal network measures when applied to neuroimaging data. This is done by applying the measures outlined above onto an fMRI dataset.

### fMRI data

Two resting-state fMRI sessions (3 Tesla, TR = 2000 ms, TE = 30 ms) from 48 healthy subjects were used in the analysis (19-31 years, 24 female). The fMRI data was downloaded from an online repository: the Beijing eyes open/eyes closed dataset available at (http://www.nitrc.org, 69). Each functional volume comprised 33 axial slices (thickness / gap= 3.5 / 0.7 mm, in-plane resolution = 64 × 64, field of view = 200 × 200 mm). The dataset contained three resting-state sessions per subject and each session lasted 480 seconds (200 image volumes, two eyes closed sessions and one eyes open session). We used data only from the 2nd or 3rd session, which were the eyes open (EO) and second eye closed session (EC), where the order was counterbalanced across subjects. Two subjects were excluded due to incomplete data. Further details regarding the scanning procedure are given in 69.

All resting state fMRI data was pre-processed using Matlab (Version 2014b, Mathworks, Inc.), CONN (70) and SPM8 (71) Matlab toolboxes. Functional imaging data was realigned and normalized to the EPI MNI template as implemented in SPM. Spatial smoothing was applied using a Gaussian filter kernel (FWHM = 8 mm). Additional image artifact regressors attributed to head movement (72, 73) were derived by using the ART toolbox for scrubbing (http://www.nitrc.org). Signal contributions from white brain matter, cerebrospinal fluid and head-movement (6 parameters), and the ART micro-movement regressors for scrubbing, were regressed out from the data using the CompCor algorithm (74, the first 5 PCA components were removed for both white matter and CSF). After regression, the data was band-passed between 0.008-0.1 Hz, as well as linearly detrended and despiked. Time series of fMRI brain activity were extracted from 264 regions of interest (spherical ROIs with a 5mm radius) using a parcellation scheme of the cortex and subcortical structures described in 11. These 264 regions of interest are further divided into 10 brain networks as described in 75.

### Creating time-graphlets (t-graphlets)

While there are many proposed methods for dynamic functional connectivity (76, 37, 69, 77, 39, 41), we chose a weighted correlation strategy (described below) as it does not require fitting any parameters or clustering.

Our logic is to calculate the dynamic functional brain connectivity estimates based on a weighted Pearson correlation. To calculate the conventional Pearson correlation coefficient, all points are weighted equally. In the weighted version points contribute differently to the coefficient depending on what weight they have been assigned. This weight is then used to calculate the weighted mean and weighted covariance to estimate the correlation coefficient. Using a unique weighting vector per time point we were able to get unique connectivity estimates for each time-point.

The weighted Pearson correlation between the signals *x* and *y* is defined as

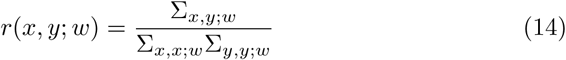

where Σ is the weighted covariance matrix and *w* is a vector of weights that is equal in length to *x* and *y*. The weighted covariance matrix is defined as

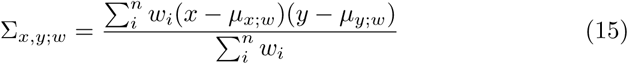

where *n* is the length of the time series. Note that Σ is the covariance matrix and 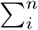 is a sum over time points. *μ*_*x;w*_ and *μ*_*y;w*_ are the weighted means, defined as

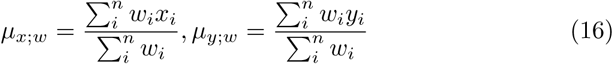

Eqs. 14-16 define the weighted Pearson coefficient with the exception of the weight vector *w*. If every element in *w* is identical, we can easily observe that the unweighted (conventional) Pearson coefficient will be calculated. Here, we instead wished to calculate a unique *w* for each time point, providing a connectivity estimate based on the weighted mean and weighted covariance.

Different weighting schemes could be applied. In fact, many of the different dynamic connectivity methods proposed in the literature are merely different weighting schemes (e.g. a non-tapered sliding window approach is just a binary weight vector). Broadly speaking, a weighting scheme between two ROIs can consider these two time series in isolation (local weighting) or, alternatively, consider every ROI’s time series (global weighting).

We decided upon the global weighting scheme, where we calculate the multivariate distance between a time-point and all other nodes. This entails that the weights for the covariance estimates at *t* are larger for other time points that display a similar global spatial pattern across all nodes to the nodes at *t*. A new weight vector is calculated for each time-point. With a unique weight vector per time-point, there is a unique weighted Pearson correlation per time-point. This reflects the weighted covariance where time-points with similar global spatial brain activation are weighted higher. This produces, for each edge, a connectivity time series with fluctuating covariance.

More formally, the weights for estimating the connectivity at time *t* are derived by taking the distance between the activation of the ROIs at *t* and each other time point (indexed by *v*):

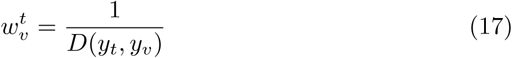

where *D* is a distance function and *y* is the multivariate time series of the ROIs. The weight vector of *t* is created by applying eq. 17 for all *v* ∈ *T,v* ≠ *t*. This implies that at the time point of interest, *t*, we calculate a vector of weights (indexed by *v*) that reflects how much the global spatial pattern of brain activity (i.e. all ROIs) differ in brain activity from *t*. For each weight vector *w^t^*, the values were scaled to range between 0 and 1. Finally, 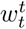 (undefined in eq. 17) is then set to 1. For the distance function, we used Euclidean distance (i.e.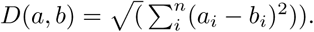

After the derivation of the connectivity time series, a Fisher transform and a Box Cox transform were applied. For the Box Cox transform the λ parameter was fit by taking maximum likelihood after a grid-search procedure through −5 and 5 in 0.1 increments for each edge. Prior to the Box Cox transformation, the smallest value was scaled to 1 to make sure the Box Cox transform performed similarly throughout the time series (78). Each connectivity time series was then standardized by subtracting the mean and dividing by the standard deviation. Binary t-graphlets were created by setting edges exceeding two standard deviations to 1, otherwise 0.

Our thresholding approach to create binary connectivity matrices is suboptimal and could be improved upon in future work (see Discussion). The need to formulate more robust thresholding practices has been an ongoing area of research in static network theory in the neurosciences (79). Similar work needs to be carried out for temporal networks as a limitation of the current approach is a heightened risk of false positive connections.

An example of a binary slice graph representation of dynamic fMRI brain connectivity is shown in Figure 5 where the connectivity within the visual subnetwork was computed using the weighted Pearson correlation method in a single subject (31 nodes belonging to the visual brain network).

**Figure 5:**
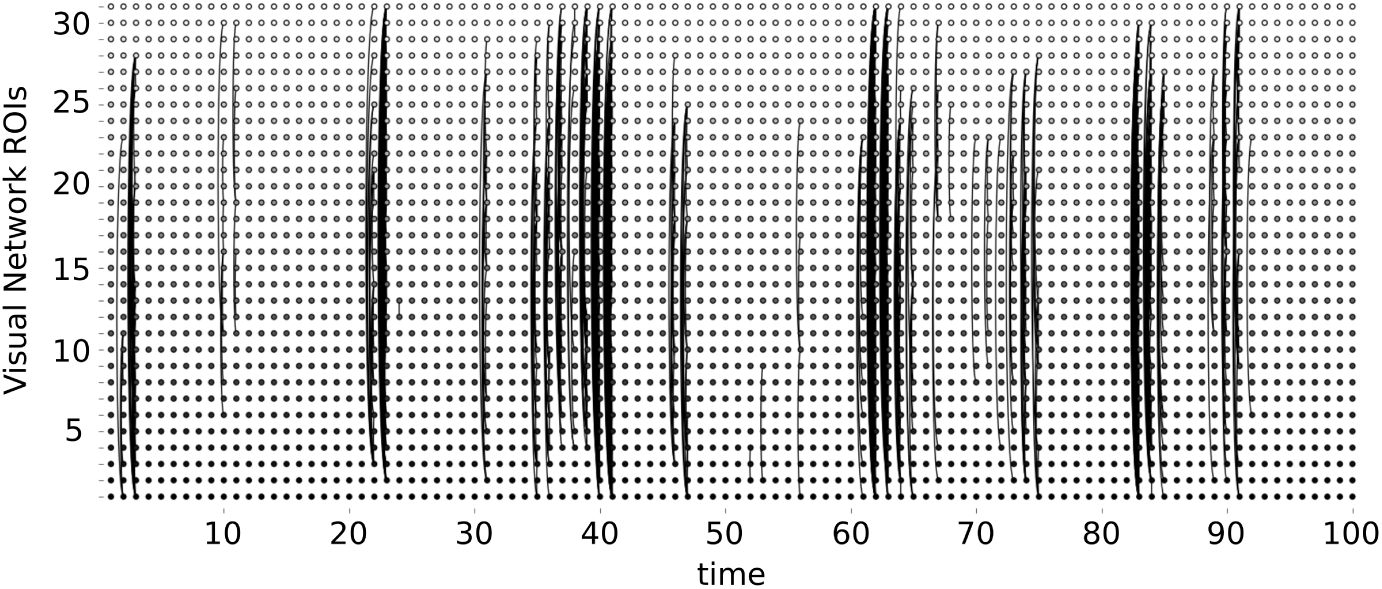
An example of the slice graph representation of temporal brain connectivity between all nodes in the visual sub-network (31 nodes, first 100 time points) computed by the use of the weighted Pearson correlation coefficient method on resting-state fMRI data (eyes open). Naturally, the visual slice graph representation becomes less interpretable as it scales up by adding more nodes.

### Tools for temporal network theory

We have implemented all temporal network measures described in the present work in a Python package of temporal network tools called *Teneto* (http://www.github.com/wiheto/teneto) for python 3.x, although the package itself is still under development. The package currently contains code for all the measures mentioned above and plotting functions for slice plots. Data formats for both the t-graphlet and contact sequence data representation are available.

### Statistics

All between group comparisons in the next section use the between group permutation method outlined previously. Null distributions were created with 100000 permutations, shuffling which group each subject’s EO/EC results belonged to and all comparisons were two tailed. For between subject comparisons a Spearman rank correlations were used.

### Applying temporal degree centrality and temporal closeness centrality

With temporal centrality measures we can formulate research questions along the following lines: (i) which nodes have the most connections through time (temporal degree centrality), or (ii) which nodes have short temporal paths to all other nodes (temporal closeness centrality). For the shortest paths calculations, we allowed all possible steps at a single time point to be used in this example.

The node centrality, averaged over subjects, was calculated for both centrality measures. The centrality estimates for nodes were compared across imaging sessions to evaluate whether there was a similar temporal pattern across subjects. The temporal degree centrality was correlated significantly for the EO and EC conditions (Figure 6A, *ρ*=0.35, p<0.0001). A similar, but slightly weaker trend was observed for temporal closeness centrality (Figure 6B, *ρ*=0.62, p<0.0001). This entails that nodes appear to have similar centrality properties for the EO and EC resting state conditions.

**Figure 6:**
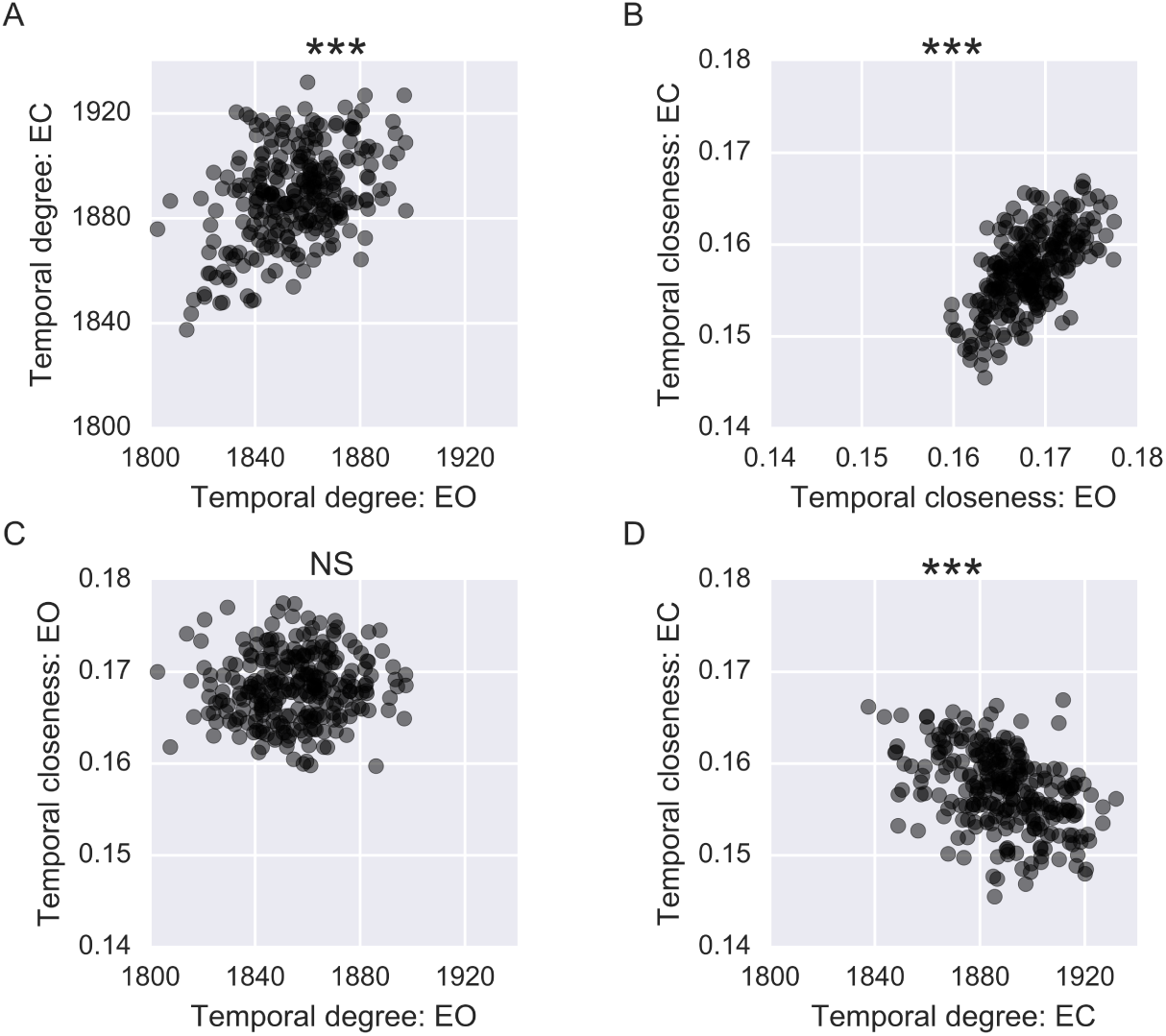
Applying temporal degree centrality and temporal closeness centrality for the EO and EC conditions. Each dot represents centrality for a node. (A) Temporal degree centrality for the EO condition versus the EC condition. (B) Same as A, but for temporal closeness centrality. (C) Temporal degree centrality versus temporal closeness centrality in the EO condition. (D) same as C, but for EC. *** signifies p<0.001

Although both centrality measures showed between session correlations, there was no consistent relationship between the two measures. The temporal degree centrality was correlated against the temporal closeness centrality. No significant relation was observed in the EO session (Figure 6C, *ρ*=0.09, p=0.15), and a negative relation for the EC session (Figure 6D, *ρ*=0.45, p<0.0001). This results suggest that the two different temporal centrality measures identify different nodes in the brain as being central.

### Applying burstiness

By applying the burstiness measure (*B*) to a fMRI dataset we can ask questions related to the temporal distribution of connections in brain connectivity. To illustrate that there is indeed a bursty pattern of brain connectivity, we first plot the distribution of all inter-contact times taken from all subjects and edges for the EO session and observe a heavy tailed distribution (Figure 7A).

**Figure 7:**
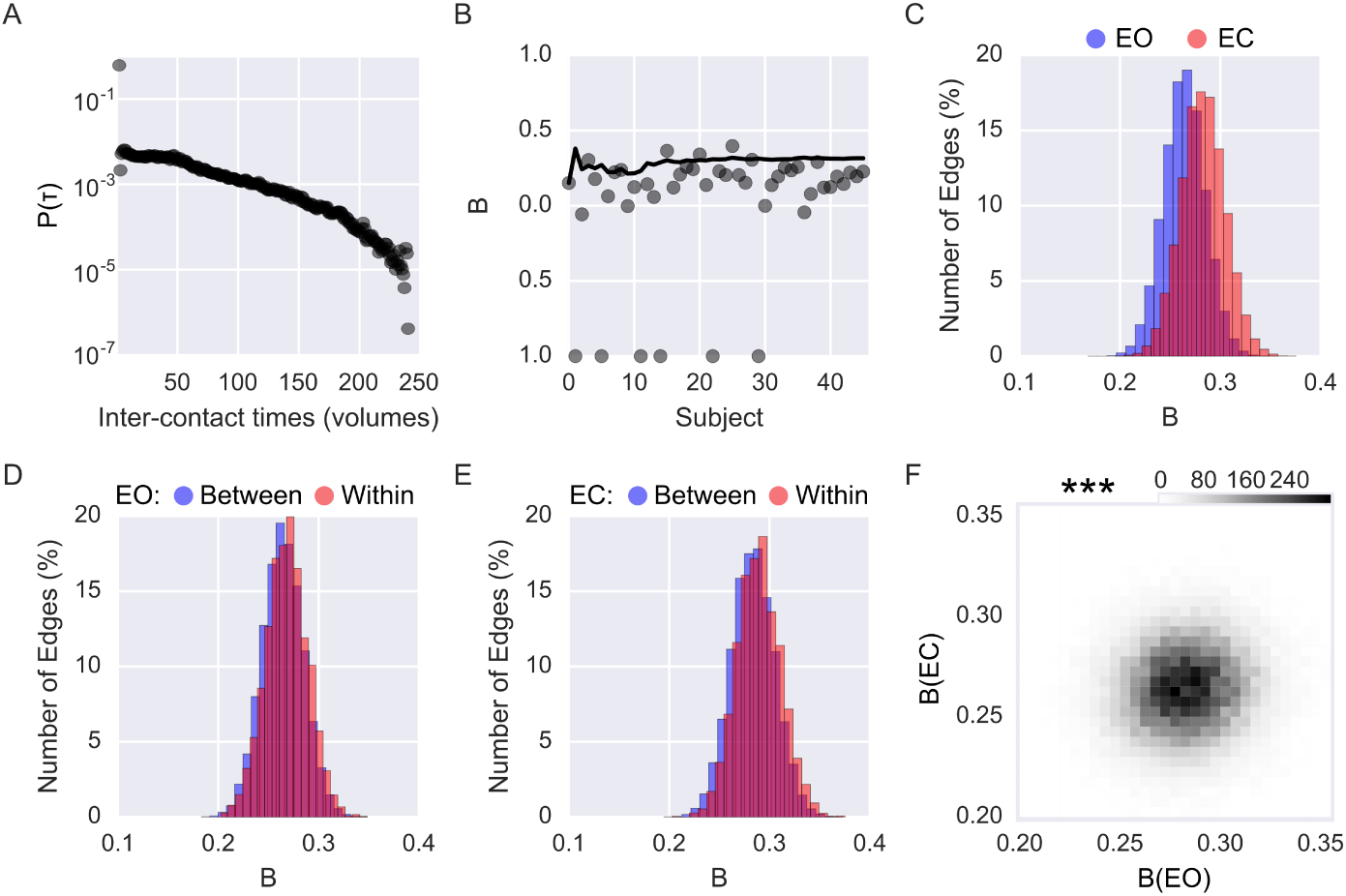
Quantifying bursty connectivity. (A) Distribution of all inter-contact probabilities, combining all edges and subjects in the eyes open condition. (B) The bursty coefficient (B) for one edge for the eyes open condition. Each dot represents *B* calculated per subject while the solid line shows the bursty coefficient when cumulatively adding subjects. Values of −1 indicate that all inter-contact times are identical (i.e. one burst, tonic connectivity or oscillations in connectivity). (C) Distribution of *B* for the different conditions (blue: EO, red: EC). (D) Same as C but showing EO within-network connectivity (red) and between-network connectivity (blue). (E) Same as D but for the EC condition. (F) Bursty coefficient for each edge across the two sessions, displayed as a heatmap. *** signifies p<0.001

We then considered the question regarding the most robust way to calculate *B*, given that our example fMRI dataset has a rather low temporal resolution and only span a limited time period. It is possible that there may not be enough edges present in each subject for a stable estimate of *B* in a single subject. To test this concern, we evaluated whether there was a difference in *B* for a single subject versus the case of concatenating the inter-contact times over multiple subjects. This was done for a single edge that connects right posterior cingulate cortex and right medial prefrontal cortex in the EO session. As shown in Figure 7B, there is a considerable variance in the individual subject estimates of burstiness. But if we cumulatively add subjects, the estimate of burstiness stabilizes after approximately 12 subjects. This illustrates the importance of having enough data to calculate reliable *B* estimates. Henceforth, all *B* estimates have been calculated by pooling inter-contact times over subjects.

We then wished to contrast EO versus EC in terms of burstiness. Both conditions showed a bursty distributions across all edges (see Figure 7C) and slightly more so for the eyes closed compared to the eyes open condition. Both within-and between-network connectivity showed a bursty distribution of connectivity patterns in both conditions (Figures 7DE).

Given that both EO and EC showed bursty correlations, we tested whether values of *B* correlated between conditions (Figure 7F). We found a weak, but significant, correlation between conditions (*ρ* = 0.066, p<0.0001). This weak correlation (i.e. less than one percent of the between-condition variance is accounted for) suggests that burstiness may relate to task related edges but more research is needed on this topic.

### Applying fluctuability

Using the fluctuability measure, researchers may ask questions regarding how many unique edges exist in a temporal network model of the dynamic functional brain connectome, indicating whether more resources (i.e. diversity of connections) are required during a given task.

The fluctuability measure was applied and contrasted between the EO and EC conditions (Figure 8A) and between-subjects (Figure 8B). We observed no significant between-subject correlation in *F* (*ρ*= 0.18, p=0.23) but found a difference for the average value of *F* between conditions (p=0.0020), with the EO condition having a higher degree of fluctuability. Thus, the EO condition may require a more varying temporal configuration of connections compared to the EC condition.

**Figure 8:**
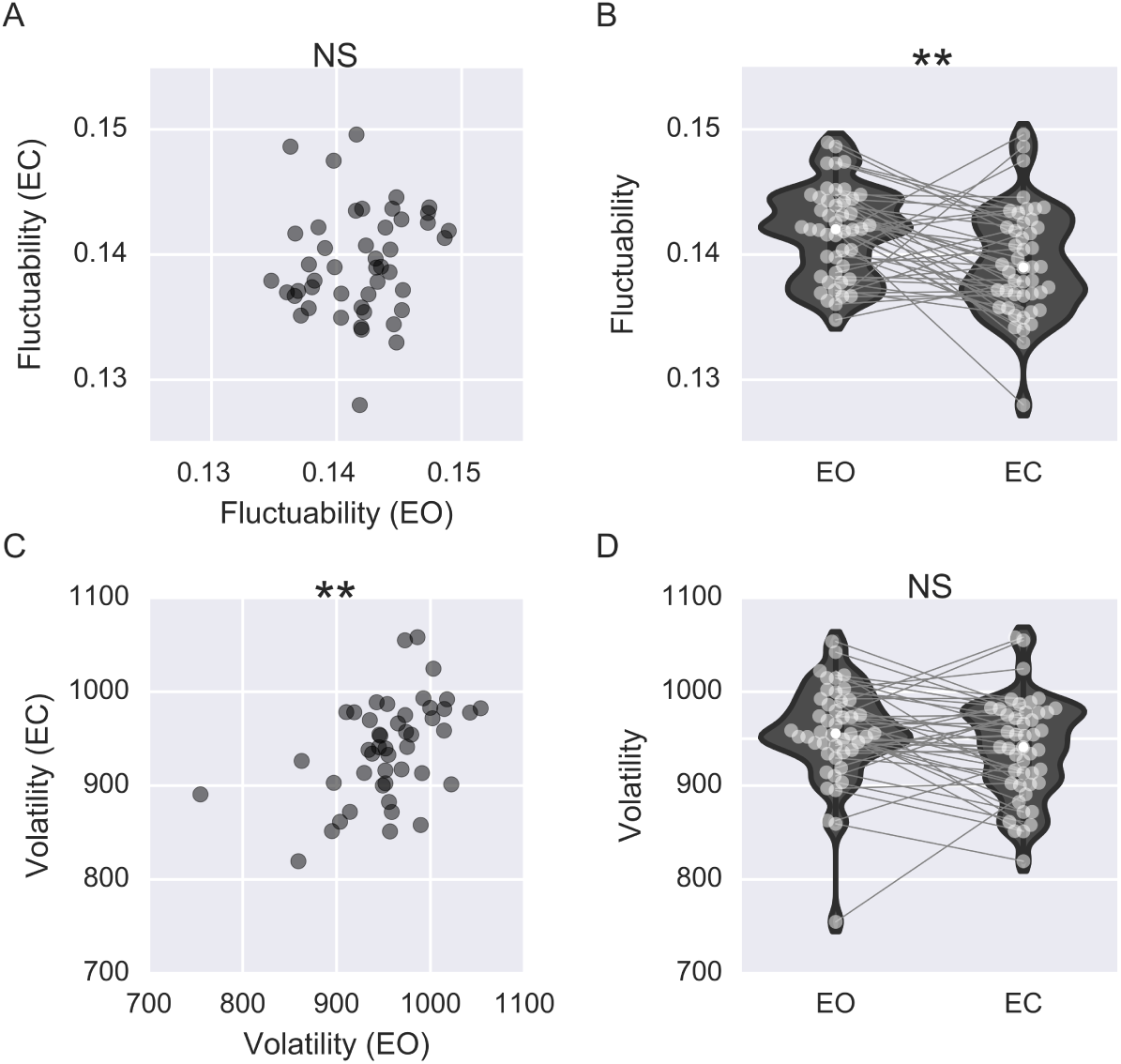
Applying fluctuability and volatility measures for the EC and EO conditions. (A) Between-subject correlation for fluctuability. (B) Violin plot showing fluctuability between the EO and EC conditions. Each light gray dot designates a subject and a line binds together data obtained from the same subjects during EO and EC conditions, respectively. For clarity, each line connecting subjects terminates at centers of the violin plots. The mean value of fluctuability for each condition is shown with a white dot. (C) Same as A but for volatility. (D) Same as B but for volatility. ** signifies p<0.01.

### Applying volatility

With volatility, we can ask whether the connectivity changes more quicker or slower through time. Some tasks might require the subject to switch between different cognitive faculties or brain states while other tasks may require the brain to be more stable and switch states less.

As with fluctuability, we computed volatility for both between-subjects (Figure 8C) and between-conditions (Figure 8D). We observed a significant correlation for between-subject volatility over the two conditions (*ρ* = 0.46, p=0.0012, Figure 8C) was obtained. Additionally, no significant difference in volatility between EO and EC was observed (p=0.051, Figure 8D).

### Applying reachability latency

The measure of reachability latency addresses the following question regarding the overall connectivity pattern along the temporal axis: how long does it take to travel to every single node in the temporal network? For example, reachability latency may be useful for evaluating the dynamics when functional or structural connectomes differ substantially. We computed the reachability latency by setting r=1 (i.e. all nodes must be reached).

The results are shown in Figure 9 and a significant difference in average reachability latency between conditions was found (Figure 9A, EO: 21.07 EC: 22.96, p=0.0005). Given that there was an overall increase in reachability latency during EC compared to EO, we decided to, post hoc, unpack this finding and check whether the discovered global difference in reachability could be localized to brain networks that should differ between the EC and EO conditions. So, rather than calculating reachability latency for the entire brain, we averaged the measure of reachability latency (to reach all nodes) for 10 preassigned brain networks (technically modules in network theory terminology). In this post hoc analysis, we see that the brain networks with the highest differences in reachability latency were the visual, dorsal attention, and the fronto-parietal brain networks (Figure 8B). Thus, the results suggests a longer reachability latency for these networks, i.e. it takes more time to reach all nodes in the visual and attention networks in the EC condition compared to EO, seems biologically reasonable.

**Figure 9:**
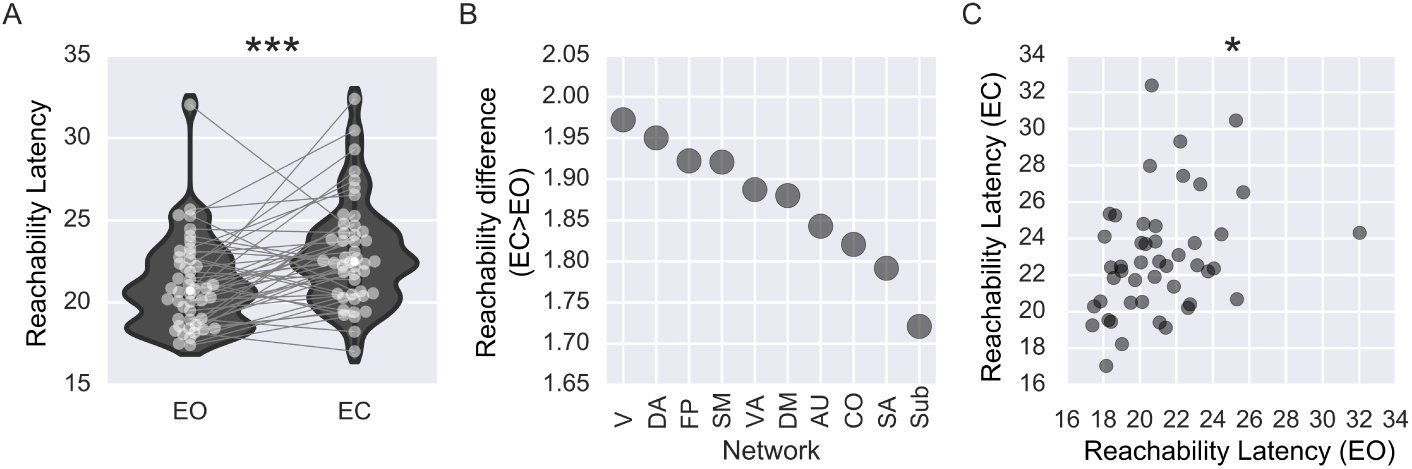
Applying reachability latency. (A) Violin plots of reachability latency for EO and EC conditions. Light gray dots correspond to a single subject and lines connect a subjects respective values between conditions. For clarity, each line connecting subjects terminates at centers of the violin plots. White dots mark the mean reachability latency. (B) Post hoc decomposition of the reachability latency difference (EC-EO) across sub-networks. (C) Between subject correlation of reachability for EO and EC. * signifies p<0.05; *** signifies p<0.001.

Despite this between-condition differences in reachability, we observed that there was also a significant between-subject relationship (*ρ*=0.36, p=0.015, Figure 8C). Taken together with the previous finding, our results show that measures of reachability latency reflects both between-conditions and between-subjects differences.

### Applying temporal efficiency

Finally, we computed the global temporal efficiency for both EO and EC conditions. While reachability latency employs the shortest temporal path to calculate how long it takes to reach a certain percentage of nodes, temporal efficiency relates to the average inverse of all shortest temporal paths.

We found that temporal efficiency is significantly larger during EO than EC (p=0.0011, Figure 10A). This finding suggests that, on average, the temporal paths are shorter in the EO condition compared to the EC condition. We observed a strong negative correlation between them during both conditions (EO: *ρ*=-0.88, p<0.0001; EC: *ρ*=-0.88, p<0.0001, see Figures 10BC). This suggests that there is a relationship between the longest shortest temporal path (part of reachability) and the average shortest temporal path (part of efficiency) in this dataset.

## Discussion

Our overarching aim in this work was to provide an overview of the key concepts of temporal networks for which we have introduced and defined temporal network measures that can be used in studies of dynamic functional brain connectivity. Additionally, we have shown the applicability of temporal metrics in network neuroscience and provided results that pertain to their characteristics by applying them onto a resting-state fMRI dataset.

### Summary of applying temporal network measures to fMRI data

Both temporal degree centrality and closeness centrality were correlated across conditions whereas no correlation between the two centrality measures was observed. This result suggests that the two centrality measures quantify different dynamic properties of the brain.

**Figure 10:**
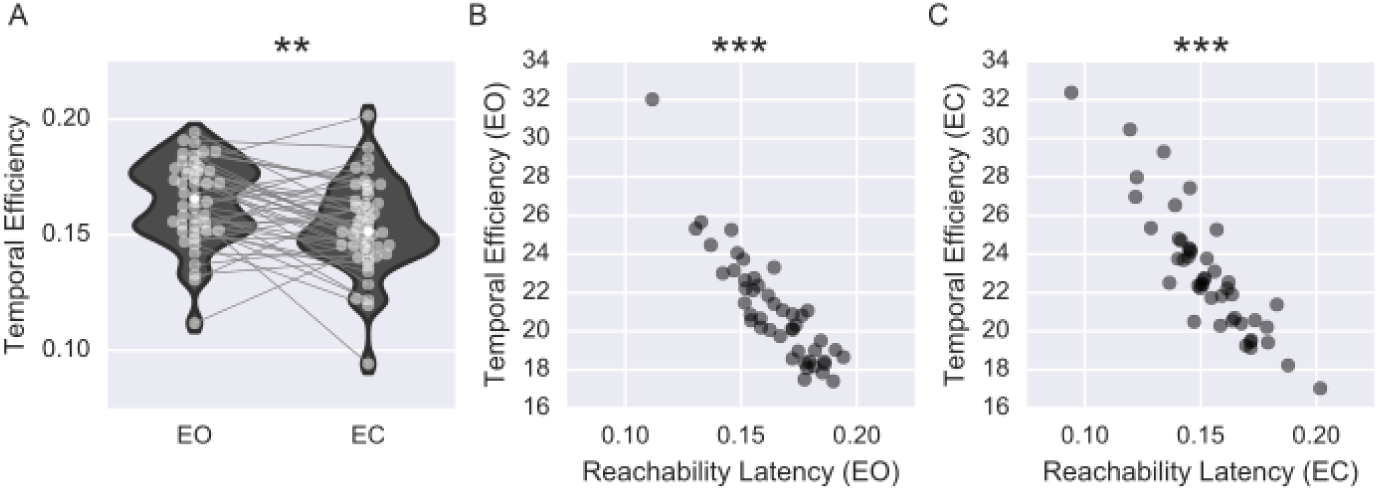
Applying global temporal efficiency and its relation with reachability latency. (A) Violin plots of global temporal efficiency for the EO and EC conditions. Light gray dots correspond to a single subject and lines connect a subjects respective values between conditions. For clarity, each line connecting subjects terminates at centers of the violin plots. White dot indicate mean global temporal efficiency. (B) Scatter plot showing each subject’s reachability latency versus temporal efficiency for the EO condition. (C) Same as B but for the EC condition. ** signifies p<0.01; *** signifies p<0.001

At a global network level, we examined the temporal uniqueness of edges (fluctuability) as well as the rate of change of connectivity (volatility). We could identify a significant condition-dependent difference in fluctuability, but no difference was observed for volatility between conditions. Conversely, a significant between-subjects correlation was found for temporal network volatility, but the between-subjects correlation in fluctuability not significant. These results suggest that the observed differences in volatility, i.e. differences in brain connectivity at two different points in time, were to a relatively larger extent driven by inter-subject differences in connectivity dynamics than by differences related to the tasks (EO/EC) per se.

Our results regarding reachability latency during EO and EC conditions indicates task driven changes in latency, especially since the connectivity of visual and attention brain networks are known to reconfigure between EO and EC conditions (80) . Thus, the observed difference in reachability latency might be a reflection of a putative network reconfiguration. Further, reachability also showed a between-subject correlation across conditions.

The distribution of inter-contact time points of connectivity between brain nodes is bursty is in agreement with our previous findings (41). Notably, our previous findings were obtained at a high temporal resolution (TR = 0.72 seconds) and it is therefore reassuring that we are able to detect similar properties of burstiness in brain connectivity also at a lower temporal resolution (TR = 2 seconds). Of note, the between-network versus within-network connectivity here differed from that obtained in a previous study which found between-network connectivity to be significantly more bursty than within-network connectivity (41). This difference is probably due to the different kind of thresholding being applied. Here a variance based thresholding was applied instead of the magnitude based in the previous study. We have discussed previously that these different strategies will prioritize different edges (81, 78).

### Other approaches to temporal network theory

The list of measures for temporal networks described here is far from exhaustive. While we have primarily focused on temporal properties that can be defined on a nodal and/or global level, detecting changes in network modularity over time is an active part of network theory research (82, 83). This has recently been applied to the brain connectome (84, 85) and applied in the context of learning (86, 87). In a similar vein, the presence of hyperedges have explored and identified groups of edges that have similar temporal evolution (88, 89). Similarly, studies investigating how different tasks evoke different network configurations (90, 75, 91) is also an active research area. Another recent exciting development is to consider a control theory based approach to network neuroscience (92), which can be applied to networks embedded in time (93).

Yet another avenue of temporal network research is to adopt static network measures to each t-graphlet and then derive time series of fluctuating static measures (94, 95). It is also possible is to quantify properties of dynamic fluctuations in brain connectivity through time and then correlate them with the underlying static network. Using such a strategy, between-subjects differences for the dynamic and the static networks can be revealed (e.g. 96).

Finally, there are considerably more measures within the temporal network literature that can be put to use within the field of network neuroscience. For example, the list of centrality measures provided here is not complete. For example, a temporal extension of the betweenness centrality, which is often used for static networks, can be adopted to the temporal domain (97). Another example is the computation of spectral centrality in the temporal domain (see 68 for further details).

### When is temporal network theory useful?

As stated in the introduction, graphs are an abstract representation corresponding to the some state in the world. The properties quantified on these representations try to reflect the properties corresponding to the world. Not every representation of brain function will require time, which makes temporal network measures unsuitable. Under what conditions will temporal network theory be of use? If the brain is viewed as a system that is processing information within different brain networks and this information is communicated between these brain networks, then we believe that temporal network theory can be advantageous to quantify these interactions.

There are a couple of additional considerations when applying temporal network theory. Interpreting what a measure means can only be done in relation to the temporal resolution of the data. For example, volatility when applied to a dataset obtained with a temporal resolution of years will obviously entail a different interpretation compared to a dataset acquired with a temporal resolution of milliseconds.

Finally, consideration is also needed about which temporal network measure(s) that should be applied to a research question. Although temporal network theory provides a wide array of measures to the users disposal, we advise against applying the entire battery of measures to a given dataset. Given a hypothesis about some state of the world (*S*), this should first be translated into a hypothesis about which network measure will quantify the network representation of *S*. A more exploratory analysis showing significant (and multiple comparison corrected) correlations in five out of ten measures, when these measures where not first formulated in relation to *S*, become hard-to-impossible to translate into something meaningful.

### Limitations and extensions for temporal network measures

Our scope was limited to temporal measures that operate on binary time series of brain connectivity (i.e. binary t-graphlets). Most of the measures discussed here can be extended and defined for series of weighted connectivity matrices. However, certain temporal measures are not straightforward to convert to the weighted case. Pertinent examples are burstiness and reachability for which no simple strategy on how to apply them in case of a weighted connectivity context is apparent.

Regardless of the method used to derive the brain connectivity time series, it is important that adequate pre-processing steps are performed on the data to avoid potential bias in the analysis. Our proposal of deriving t-graphlets with weighted Pearson correlation coefficients to compute time series of brain connectivity constitute no exception to this concern. In a connectivity analysis that is based on sequences of binary t-graphlets, an absence or presence of an edge might potentially be influenced by the user’s selection of thresholding. Hence, the strategy regarding how to optimally threshold the t-graphles into binary graphlets is of vital importance. We believe that it is important to keep in mind that comparisons of variance as well as the mean of connectivity time series might be biased by the underlying mean-variance relationship (81, 78). This further emphasizes the need for adequate thresholding strategies for connectivity time series. Moreover, subject head motion, known to be a large problem for fMRI connectivity studies, (72, 73, 98), can also lead to spurious dynamic properties (99) and should be controlled for.

### Outlook

By providing a survey of the theory of temporal networks and by showing their applicability and usefulness in network neuroscience, we hope that we have stirred the readers interest in using models based on temporal networks when studying the dynamics of functional brain connectivity. To this end, we have implemented all temporal network measures described in the present paper in a software package that is freely available ( *Teneto*, written in Python and can be downloaded at github.com/wiheto/teneto). The plan for the *Teneto* package is to include additional temporal network measures, plotting routines, wrappers for other programming languages, and dynamic connectivity estimation.

## Acknowledgements

We would like to thank Pontus Plavén-Sigray, Björn Schiffler, Granville Matheson, Simon Skau and Lieke de Boer for helpful comments and discussions about the manuscript.

## Funding

P.F. was supported by the Swedish Research Council (grant no. 621-2012-4911) and the Swedish e-Science Research Center.

## Author contributions

WHT, PB, PF designed the study. WHT wrote all code. WHT and PF wrote the paper. WHT, PB, PF revised the paper.

## References

[1] O. Sporns, G. Tononi, and R. Kötter, “The human connectome: A structural description of the human brain,” PLoS Computational Biology, vol. 1, no. 4, pp. 0245–0251, 2005.

[2] O. Sporns, “Networks of the Brain,” MIT press, Cambridge, MA, 2009.

[3] M. D. Greicius, B. Krasnow, A. L. Reiss, and V. Menon, “Functional connectivity in the resting brain: a network analysis of the default mode hypothesis,” Proc.Natl.Acad.Sci.U.S A, vol. 100, nos. 0027-8424 (Print) LA - eng PT - Journal Article PT - Research Support, Non-U.S. Gov’t PT - Research Support, U.S. Gov’t, P.H.S SB - IM, pp. 253–258, 2003.

[4] P. Fransson, “Spontaneous low-frequency BOLD signal fluctuations: An fMRI investigation of the resting-state default mode of brain function hypothesis,” Human Brain Mapping, vol. 26, no. 1, pp. 15–29, Sep. 2005.

[5] M. D. Fox, A. Z. Snyder, J. L. Vincent, M. Corbetta, D. C. Van Essen, and M. E. Raichle, “The human brain is intrinsically organized into dynamic, anticorrelated functional networks.” Proceedings of the National Academy of Sciences of the United States of America, vol. 102, no. 27, pp. 9673–8, Jul. 2005.

[6] S. M. Smith, “Correspondence of the brain’s functional architecture during activation and rest.” Proceedings of the National Academy of Sciences of the

[7] F. de Pasquale, “Temporal dynamics of spontaneous MEG activity in brain networks.” Proceedings of the National Academy of Sciences of the United States of America, vol. 107, no. 13, pp. 6040–5, Mar. 2010.

[8] M. J. Brookes, “Investigating the electrophysiological basis of resting state networks using magnetoencephalography.” Proceedings of the National Academy of Sciences of the United States of America, vol. 108, no. 40, pp. 16783–16788, 2011.

[9] J. F. Hipp, D. J. Hawellek, M. Corbetta, M. Siegel, and A. K. Engel, “Large-scale cortical correlation structure of spontaneous oscillatory activity,” Nature Neuroscience, vol. 15, no. 6, pp. 884–890, 2012.

[10] R. L. Buckner, “Cortical hubs revealed by intrinsic functional connectivity: mapping, assessment of stability, and relation to Alzheimer’s disease.” The Journal of neuroscience: the official journal of the Society for Neuroscience, vol. 29, no. 6, pp. 1860–73, Mar. 2009.

[11] J. D. Power, “Functional Network Organization of the Human Brain,” Neuron, vol. 72, no. 4, pp. 665–678, Nov. 2011.

[12] J. D. Power, B. L. Schlaggar, C. N. Lessov-Schlaggar, and S. E. Petersen, “Evidence for hubs in human functional brain networks.” Neuron, vol. 79, no. 4, pp. 798–813, Aug. 2013.

[13] E. H. J. Nijhuis, A.-M. van Cappellen van Walsum, and D. G. Norris, “Topographic hub maps of the human structural neocortical network.” PloS one, vol. 8, no. 6, p. e65511, Jan. 2013.

[14] M. D. Fox and M. Greicius, “Clinical applications of resting state functional connectivity.” Frontiers in systems neuroscience, vol. 4, no. June, p. 19, Jan. 2010.

[15] D. Zhang and M. E. Raichle, “Disease and the brain’s dark energy.” Nature reviews. Neurology, vol. 6, no. 1, pp. 15–28, Jan. 2010.

[16] G. Deco, V. K. Jirsa, and A. R. McIntosh, “Emerging concepts for the dynamical organization of resting-state activity in the brain.” Nature reviews. Neuroscience, vol. 12, no. 1, pp. 43–56, Jan. 2011.

[17] J. Kelso, Dynamic Patterns: The Self-organization of Brain and Behavior. 1995.

[18] E. Tognoli and J. a S. Kelso, “Brain coordination dynamics: true and false faces of phase synchrony and metastability.” Progress in neurobiology, vol. 87, no. 1, pp. 31–40, Jan. 2009.

[19] E. Tognoli and J. a S. Kelso, “The metastable brain.” Neuron, vol. 81, no. 1, pp. 35–48, Jan. 2014.

[20] S. L. Bressler and J. a.S. Kelso, “Cortical coordination dynamics and cognition,” Trends in cognitive sciences, vol. 5, no. 1, pp. 26–36, Jan. 2001.

[21] G. Buzsáki and A. Draguhn, “Neuronal oscillations in cortical networks.” Science (New York, N.Y.), vol. 304, no. 5679, pp. 1926–1929, Jun. 2004.

[22] G. Buzsáki, Rhythms of the Brain. Oxford University Press, 2006.

[23] M. Siegel, T. H. Donner, and A. K. Engel, “Spectral fingerprints of large-scale neuronal interactions.” Nature reviews. Neuroscience, vol. 13, no. 2, pp. 121–34, Feb. 2012.

[24] G. Buzsáki, “Theta rhythm of navigation: Link between path integration and landmark navigation, episodic and semantic memory,” Hippocampus, vol. 15, no. 7, pp. 827–840, 2005.

[25] S. M. Montgomery and G. Buzsáki, “Gamma oscillations dynamically couple hippocampal CA3 and CA1 regions during memory task performance.” Proceedings of the National Academy of Sciences of the United States of America, vol. 104, no. 36, pp. 14495–14500, 2007.

[26] U. Friese, M. Köster, U. Hassler, U. Martens, N. Trujillo-Barreto, and T. Gruber, “Successful memory encoding is associated with increased cross-frequency coupling between frontal theta and posterior gamma oscillations in human scalp-recorded EEG,” NeuroImage, vol. 66, pp. 642–647, 2013.

[27] P. Fries, “A mechanism for cognitive dynamics: neuronal communication through neuronal coherence.” Trends in cognitive sciences, vol. 9, no. 10, pp. 474–80, Oct. 2005.

[28] C. a Bosman, “Attentional stimulus selection through selective synchronization between monkey visual areas.” Neuron, vol. 75, no. 5, pp. 875–88, Sep. 2012.

[29] S. L. Bressler, C. G. Richter, Y. Chen, and M. Ding, “Cortical functional network organization from autoregressive modeling of local field potential oscillations,” Statistics in Medicine, vol. 26, no. 21, pp. 3875–3885, Sep. 2007.

[30] T. J. Buschman and E. K. Miller, “Top-down versus bottom-up control of attention in the prefrontal and posterior parietal cortices.” Science (New York, N.Y.), vol. 315, no. 5820, pp. 1860–1862, 2007.

[31] A. M. Bastos, W. M. Usrey, R. A. Adams, G. R. Mangun, P. Fries, and K. J. Friston, “Canonical Microcircuits for Predictive Coding,” Neuron, vol. 76, no. 4, pp. 695–711, Nov. 2012.

[32] A. M. Bastos, “Visual areas exert feedforward and feedback influences through distinct frequency channels,” Neuron, vol. 85, no. 2, pp. 390–401, 2015.

[33] C. G. Richter, W. H. Thompson, C. A. Bosman, and P. Fries, “Top-down modulation of stimulus drive via beta-gamma cross-frequency interaction.” May 2016.

[34] C. G. Richter, R. Coppola, and S. L. Bressler, “Top-Down Beta Oscillatory Signaling Conveys Behavioral Context to Primary Visual Cortex,” bioRxiv, p. 074609, 2016.

[35] G. Michalareas, J. Vezoli, S. van Pelt, J. M. Schoffelen, H. Kennedy, and P. Fries, “Alpha-Beta and Gamma Rhythms Subserve Feedback and Feedforward Influences among Human Visual Cortical Areas,” Neuron, vol. 89, no. 2, pp. 384–397, 2016.

[36] T. van Kerkoerle, “Alpha and gamma oscillations characterize feedback and feedforward processing in monkey visual cortex,” Proceedings of the National Academy of Sciences, vol. 111, no. 40, pp. 14332–41, 2014.

[37] E. a Allen, E. Damaraju, S. M. Plis, E. B. Erhardt, T. Eichele, and V. D. Calhoun, “Tracking whole-brain connectivity dynamics in the resting state.” Cerebral cortex (New York, N.Y.: 1991), vol. 24, no. 3, pp. 663–76, Mar. 2014.

[38] R. M. Hutchison, “Dynamic functional connectivity: Promise, issues, and interpretations,” NeuroImage, vol. 80, pp. 360–378, Oct. 2013.

[39] J. M. Shine, O. Koyejo, P. T. Bell, K. J. Gorgolewski, M. Gilat, and R. A. Poldrack, “Estimation of dynamic functional connectivity using Multiplication of Temporal Derivatives,” NeuroImage, vol. 122, pp. 399–407, 2015.

[40] W. H. Thompson and P. Fransson, “The frequency dimension of fMRI dynamic connectivity: network connectivity, functional hubs and integration in the resting brain,” NeuroImage, vol. 121, pp. 227–242, 2015.

[41] W. H. Thompson and P. Fransson, “Bursty properties revealed in large-scale brain networks with a point-based method for dynamic functional connectivity,” Scientific Reports, vol. 6, no. November, p. 39156, Dec. 2016.

[42] J. M. Shine, O. Koyejo, and R. A. Poldrack, “Temporal meta-states are associated with differential patterns of dynamic connectivity, network topology and attention,” no. 10, pp. 1–4, 2016.

[43] F. de Pasquale, “A cortical core for dynamic integration of functional networks in the resting human brain.” Neuron, vol. 74, no. 4, pp. 753–64, May 2012.

[44] A. P. Baker, “Fast transient networks in spontaneous human brain activity,” eLife, vol. 2014, no. 3, pp. 1–18, 2014.

[45] N. J. Kopell, H. J. Gritton, M. A. Whittington, and M. A. Kramer, “Beyond the connectome: The dynome,” Neuron, vol. 83, no. 6, pp. 1319–1328, 2014.

[46] V. D. Calhoun, R. Miller, G. Pearlson, and T. Adalı, “The Chronnectome: Time-Varying Connectivity Networks as the Next Frontier in fMRI Data Discovery,” Neuron, vol. 84, no. 2, pp. 262–274, Oct. 2014.

[47] M. Newman, Networks. An introduction. 2010, p. 772.

[48] E. Bullmore and O. Sporns, “Complex brain networks: graph theoretical analysis of structural and functional systems.” Nature reviews. Neuroscience, vol. 10, no. 3, pp. 186–98, Mar. 2009.

[49] M. Berlingerio, M. Coscia, F. Giannotti, A. Monreale, and D. Pedreschi, “Foundations of multidimensional network analysis,” Proceedings - 2011 International Conference on Advances in Social Networks Analysis and Mining, ASONAM 2011, pp. 485–489, 2011.

[50] P. Basu, A. Bar-noy, M. P. Johnson, and R. Ramanathan, “Modeling and Analysis of Time-Varying Graphs,” arXiv preprint arXiv:1012.0260., 2010.

[51] P. Holme and J. Saramäki, “Temporal networks,” Physics Reports, vol. 519, no. 3, pp. 97–125, 2012.

[52] R. K. Pan and J. Saramäki, “Path lengths, correlations, and centrality in temporal networks,” Physical Review E - Statistical, Nonlinear, and Soft Matter Physics, vol. 84, no. 1, 2011.

[53] A.-L. Barabási, “The origin of bursts and heavy tails in human dynamics,” vol. Nature, no. 435, pp. 207–211, May 2005.

[54] A. Vázquez, J. G. Oliveira, Z. Dezsö, K. I. Goh, I. Kondor, and A. L. Barabási, “Modeling bursts and heavy tails in human dynamics,” Physical Review E - Statistical, Nonlinear, and Soft Matter Physics, vol. 73, no. 3, pp. 1–19, 2006.

[55] A. Vazquez, B. Rácz, A. Lukács, and A. L. Barabási, “Impact of non-poissonian activity patterns on spreading processes,” Physical Review Letters, vol. 98, no. 15, pp. 1–4, 2007.

[56] A.-L. Barabási, Bursts: The Hidden Pattern Behind Everything We Do. Penguin, 2010.

[57] B. Min, K.-I. Goh, and A. Vazquez, “Spreading dynamics following bursty human activity patterns,” Physical Review E, vol. 83, no. 3, p. 036102, Mar. 2011.

[58] J.-P. Eckmann, E. Moses, and D. Sergi, “Entropy of dialogues creates coherent structures in e-mail traffic.” Proceedings of the National Academy of Sciences of the United States of America, vol. 101, no. 7, pp. 14333–14337, 2004.

[59] H. H. Jo, M. Karsai, J. Kertész, and K. Kaski, “Circadian pattern and burstiness in human communication activity,” New Journal of Physics, vol. 14, p. 013055, 2012.

[60] A. Vazquez, “Spreading Dynamics Following Bursty Activity Patterns,” in Temporal networks, Springer, 2013, pp. 161–174.

[61] L. E. C. Rocha, F. Liljeros, and P. Holme, “Information dynamics shape the sexual networks of Internet-mediated prostitution,” Proceedings of the National Academy of Sciences, vol. 107, no. 13, pp. 5706–5711, Mar. 2010.

[62] T. Takaguchi, N. Masuda, and P. Holme, “Bursty Communication Patterns Facilitate Spreading in a Threshold-Based Epidemic Dynamics,” PloS one, vol. 8, no. 7, p. 096461, 2013.

[63] F. Freyer, K. Aquino, P. a Robinson, P. Ritter, and M. Breakspear, “Bistability and non-Gaussian fluctuations in spontaneous cortical activity.” The Journal of neuroscience: the official journal of the Society for Neuroscience, vol. 29, no. 26, pp. 8512–8524, 2009.

[64] F. Freyer, J. a. Roberts, P. Ritter, and M. Breakspear, “A Canonical Model of Multistability and Scale-Invariance in Biological Systems,” PLoS Computational Biology, vol. 8, no. 8, 2012.

[65] J. a Roberts, T. W. Boonstra, and M. Breakspear, “The heavy tail of the human brain,” Current Opinion in Neurobiology, vol. 31, pp. 164–172, 2015.

[66] K.-I. Goh and a.-L. Barabási, “Burstiness and memory in complex systems,” EPL (Europhysics Letters), vol. 81, no. 4, p. 48002, Feb. 2008.

[67] P. Holme, “Network reachability of real-world contact sequences,” Physical Review E - Statistical, Nonlinear, and Soft Matter Physics, vol. 71, no. 4, pp. 1–8, 2005.

[68] V. Nicosia, J. Tang, C. Mascolo, M. Musolesi, G. Russo, and V. Latora, “Graph Metrics for Temporal Networks,” in Temporal networks, Berlin Heidelberg: Springer, 2013, pp. 15–40.

[69] X. Liu and J. H. Duyn, “Time-varying functional network information extracted from brief instances of spontaneous brain activity.” Proceedings of the National Academy of Sciences of the United States of America, vol. 110, no. 11, pp. 4392–7, Mar. 2013.

[70] S. Whitfield-Gabrieli and A. Nieto-Castanon, “Conn: A Functional Connectivity Toolbox for Correlated and Anticorrelated Brain Networks,” Brain Connectivity, vol. 2, no. 3, pp. 125–141, 2012.

[71] K. J. Friston, a. P. Holmes, K. J. Worsley, J.-P. Poline, C. D. Frith, and R. S. J. Frackowiak, “Statistical parametric maps in functional imaging: A general linear approach,” Human Brain Mapping, vol. 2, no. 4, pp. 189–210, 1995.

[72] K. R. A. van Dijk, M. R. Sabuncu, and R. L. Buckner, “The influence of head motion on intrinsic functional connectivity MRI,” NeuroImage, vol. 59, no. 1, pp. 431–438, Jan. 2012.

[73] J. D. Power, K. a Barnes, A. Z. Snyder, B. L. Schlaggar, and S. E. Petersen, “Spurious but systematic correlations in functional connectivity MRI networks arise from subject motion.” NeuroImage, vol. 59, no. 3, pp. 2142–54, Feb. 2012.

[74] Y. Behzadi, K. Restom, J. Liau, and T. T. Liu, “A component based noise correction method (CompCor) for BOLD and perfusion based fMRI,” NeuroImage, vol. 37, no. 1, pp. 90–101, 2007.

[75] M. W. Cole, J. R. Reynolds, J. D. Power, G. Repovs, A. Anticevic, and T. S. Braver, “Multi-task connectivity reveals flexible hubs for adaptive task control.” Nature neuroscience, vol. 16, no. 9, pp. 1348–55, Sep. 2013.

[76] S. M. Smith, “Temporally-independent functional modes of spontaneous brain activity.” Proceedings of the National Academy of Sciences of the United States of America, vol. 109, no. 8, pp. 3131–6, Feb. 2012.

[77] M. A. Lindquist, Y. Xu, M. B. Nebel, and B. S. Caffo, “Evaluating dynamic bivariate correlations in resting-state fMRI: A comparison study and a new approach,” NeuroImage, vol. 101, pp. 531–546, 2014.

[78] W. H. Thompson and P. Fransson, “On Stabilizing the Variance of Dynamic Functional Brain Connectivity Time Series,” Brain Connectivity, vol. 6, no. 10, pp. 735–746, Dec. 2016.

[79] M. Drakesmith, K. Caeyenberghs, a. Dutt, G. Lewis, a.S. David, and D. Jones, “Overcoming the effects of false positives and threshold bias in graph theoretical analyses of neuroimaging data,” NeuroImage, vol. 118, pp. 313–333, 2015.

[80] D. Zhang, “Directionality of large-scale resting-state brain networks during eyes open and eyes closed conditions,” Frontiers in Human Neuroscience, vol. 9, no. February, p. Article 81, 2015.

[81] W. H. Thompson and P. Fransson, “The mean–variance relationship reveals two possible strategies for dynamic brain connectivity analysis in fMRI,” Frontiers in Human Neuroscience, vol. 9, no. July, pp. 1–7, 2015.

[82] P. J. Mucha, T. Richardson, K. Macon, M. a Porter, and J.-P. Onnela, “Community structure in time-dependent, multiscale, and multiplex networks.” Science (New York, N.Y.), vol. 328, no. 5980, pp. 876–8, May 2010.

[83] M. Rosvall and C. T. Bergstrom, “Mapping change in large networks.” PloS one, vol. 5, no. 1, p. e8694, Jan. 2010.

[84] A. V. Mantzaris, “Dynamic network centrality summarizes learning in the human brain,” Journal of Complex Networks, vol. 1, no. 1, pp. 83–92, 2013.

[85] D. S. Bassett, M. A. Porter, N. F. Wymbs, S. T. Grafton, J. M. Carlson, and P. J. Mucha, “Robust detection of dynamic community structure in networks,” Chaos, vol. 23, no. 1, 2013.

[86] D. S. Bassett, N. F. Wymbs, M. a Porter, P. J. Mucha, J. M. Carlson, and S. T. Grafton, “Dynamic reconfiguration of human brain networks during learning.” Proceedings of the National Academy of Sciences of the United States of America, vol. 108, no. 18, pp. 7641–6, May 2011.

[87] D. S. Bassett, M. Yang, N. F. Wymbs, and S. T. Grafton, “Learning-Induced Autonomy of Sensorimotor Systems,” Nature neuroscience, vol. 18, no. 5, pp. 744–751, 2015.

[89] E. N. Davison, “Brain Network Adaptability across Task States,” PLoS Computational Biology, vol. 11, no. 1, 2015.

[89] E. N. Davison, “Individual Differences in Dynamic Functional Brain Connectivity Across the Human Lifespan,” arXiv preprint, pp. 1–26, 2016.

[90] M. Ekman, J. Derrfuss, M. Tittgemeyer, and C. J. Fiebach, “Predicting errors from reconfiguration patterns in human brain networks,” Proceedings of the National Academy of Sciences of the United States of America, vol. 109, no. 41, pp. 16714–16719, 2012.

[91] M. G. Mattar, M. W. Cole, and L. Sharon, “A Functional Cartography of Cognitive Systems,” pp. 1–36, 2014.

[92] S. Gu, “Controllability of structural brain networks,” Nature Communications, vol. 6, p. 8414, 2015.

[93] S. Gu, “Optimal Trajectories of Brain State Transitions,” pp. 1–10, 2016.

[94] M. Bola and B. A. Sabel, “Dynamic reorganization of brain functional networks during cognition,” NeuroImage, vol. 114, pp. 398–413, 2015.

[95] S. Chiang, “Time-dependence of graph theory metrics in functional connectivity analysis,” NeuroImage, vol. 125, pp. 601–615, 2016.

[96] R. F. Betzel, M. Fukushima, Y. He, X. N. Zuo, and O. Sporns, “Dynamic fluctuations coincide with periods of high and low modularity in resting-state functional brain networks,” NeuroImage, vol. 127, no. February 2016, pp. 287–297, 2016.

[97] J. Tang, M. Musolesi, C. Mascolo, V. Latora, and V. Nicosia, “Analysing Information Flows and Key Mediators through Temporal Centrality Metrics Categories and Subject Descriptors,” Proceedings of the 3rd Workshop on Social Network Systems, no. Figure 1, p. 3, 2010.

[98] J. D. Power, B. L. Schlaggar, and S. E. Petersen, “Recent progress and outstanding issues in motion correction in resting state fMRI,” NeuroImage, vol. 105, pp. 536–551, 2015.

[99] T. O. Laumann, “On the Stability of BOLD fMRI Correlations,” Cerebral Cortex, pp. 1–14, 2016.

